# Exploring Population Differences in the Human Gut Microbiome: from Microbial Abundance to Single Nucleotide Polymorphisms

**DOI:** 10.1101/2025.04.10.648293

**Authors:** Jingjing Wang, Zhenmiao Zhang, Yang Chen, Xi Zhou, Jiaxin Xiang, Chao Yang, Dmitry A. Rodionov, Andrei L. Osterman, Qinwei Qiu, Yusheng Deng, Yanmin Liu, Chengrui Wang, Xiaoxiao Shang, Li Huang, Chen Sun, Jianwen Guo, Zhimin Yang, Lijuan Han, Lixiang Zhai, Zhaoxiang Bian, Wei Jia, Xiaodong Fang, Lu Zhang

**Affiliations:** Department of Computer Science, Hong Kong Baptist University, Street, Hong Kong SAR, China; State Key Laboratory of Traditional Chinese Medicine Syndrome, The Second Affiliated Hospital of Guangzhou University of Chinese, Guangzhou, China; State Key Laboratory of Dampness Syndrome of Chinese Medicine, The Second Affiliated Hospital of Guangzhou University of Chinese, Guangzhou, China; Sanford Burnham Prebys Medical Discovery Institute, La Jolla, CA, United States; Department of Scientific Research, Kangmeihuada GeneTech Co., Ltd (KMHD), Shenzhen, China; Centre for Chinese Herbal Medicine Drug Development Limited, Hong Kong Baptist University, Hong Kong SAR, China; School of Chinese Medicine, Hong Kong Baptist University, Hong Kong SAR, China; Department of Pharmacology and Pharmacy, The University of Hong Kong, Hong Kong SAR, China; Institute for Research and Continuing Education, Hong Kong Baptist University, Shenzhen, China

**Keywords:** human gut microbiome, single nucleotide polymorphisms, microbial reference genomes, population differences

## Abstract

The human gut microbiota exhibits significant diversity across populations, influenced by factors such as geography, diet, and lifestyle, particularly between the Han Chinese and non-Chinese populations. While previous studies have predominantly focused on the taxonomic abundance of the gut microbiome, the impact of single nucleotide polymorphisms (SNPs) in driving population-specific differences remains largely underexplored.

In this study, we systematically investigated gut microbial differences between the Han Chinese and non-Chinese populations using the Human Gut Microbiome Reference Genome Catalog (HGMRGC). HGMRGC includes 271,480 non-redundant genomes from 5,785 prokaryotic species, which was generated by metagenomic sequencing data from 9,320 publicly available samples across 22 countries and 3,584 newly sequenced samples from Han Chinese individuals across 29 provinces and regions in China.

We observed geography was the primary driver of microbial variation of abundance and SNPs. We identified 625 novel population-specific genome clusters from HGMRGC with functional differences in carbohydrate utilization and 126 species exhibiting distinct prevalence related to vitamin biosynthesis, antibiotic resistance, and carbohydrate metabolism. Beta diversity analysis highlighted significant inter-population differences in both microbial abundance and SNPs, while alpha diversity analysis revealed that non-Chinese populations exhibited higher diversity in microbial abundance, and Han Chinese populations displayed greater diversity in SNPs. These results provide valuable insights into population-specific microbial diversity, laying the groundwork for future research on its functional and health implications.

## Introduction

The human gastrointestinal tract harbors one of the most diverse and complex microbial ecosystems in the body [1], playing an essential role in maintaining health and modulating diseases [2–5]. Advances in high-throughput sequencing technologies have substantially reduced the cost of metagenomic sequencing, enabling systematic investigations of the human gut microbiota and their variations across populations [6]. Geography and ethnicity have been proven as key factors shaping gut microbiota composition. Previous studies have revealed striking regional differences in microbial taxa [7, 8]. For example, *Prevotella* is more abundant in non-Western populations [7], while *Bacteroides* is more common in Han Chinese [8]. However, most of these studies focus on taxonomic composition and microbial relative abundances, often neglecting the effect of single nucleotide polymorphisms (SNPs), which play a pivotal role in shaping microbial metabolic functions and characteristics [9].

Recent advances in microbial culturomics [10–12] and metagenome-assembled genomes (MAGs) from metagenome assembly [13–15] have expanded our understanding of microbial diversity, enabling the construction of gut microbial reference catalogs to explore microbial SNPs. Several studies [9, 16, 17] have explored the links between microbial SNPs and host traits using available human gut reference genomes. For example, SNPs in *Bilophila wadsworthia* were associated with host dietary patterns and lipopolysaccharide abundance, potentially influencing weight gain [9]. A high-fiber diet intervention notably altered the gut microbiota composition, resulting in increased SNP proportions of genes related to amino acid biosynthetic in *Faecalibacterium prausnitzii* among children with Prader–Willi syndrome [17]. While these studies emphasize the relationship between microbial SNPS and host traits, the role of microbial SNPs in driving population-specific differences among ethnic groups remains largely unexplored. This is primarily due to the bias of current reference genome catalogs toward Western populations [18], which limits their ability to support studies examining differences in microbial genomic variations among diverse populations. For example, only 22% of the samples in the Unified Human Gastrointestinal Genome (UHGG) version 2 [19], the most extensive reference collection, are sourced from Chinese populations. Furthermore, the existing catalogs [13– 15, 19, 20] rely on publicly available datasets that lack detailed metadata to ensure thorough inclusion of diverse geographical regions and population groups, particularly for the Han Chinese population, which is marked by significant dietary and lifestyle diversity across regions. Addressing these biases is critical for improving the completeness and applicability of gut microbial reference genomes.

In this study, we thoroughly investigated gut microbial differences between the Han Chinese (HC) and non-Chinese (NC) populations based on both microbial abundance and SNPs using a high-quality Human Gut Microbiome Reference Genome Catalog (HGMRGC). The HGMRGC comprises 271,480 non-redundant genomes representing 5,785 prokaryotic species, offering significant improvements over existing resources. HGMRGC was generated using metagenomic sequencing data from 9,320 public-available samples across 22 countries and 3,584 newly sequenced Han Chinese samples across 29 provinces and regions in China. Our analyses revealed geography as the primary driver of microbial variation, surpassing age, sex, and body mass index (BMI). We identified 625 novel population-specific genome clusters in HGMRGC, which were observed to be involved in different functions in carbohydrate utilization, and 126 species with distinct prevalence linked to vitamin biosynthesis, antibiotic resistance, and carbohydrate metabolism. Additionally, beta diversity analysis highlighted significant inter-population differences in both microbial abundance and SNPs, while alpha diversity analysis showed NC populations exhibited higher diversity in microbial abundance, and HC populations showed greater diversity in microbial SNPs. These findings provide valuable insights into population-specific microbial diversity, laying the groundwork for future research on its functional and health implications.

## Results

### Construct a high-quality catalog of human gut microbial reference genomes

We developed a high-quality Human Gut Microbial Reference Genome Catalog (HGMRGC) by integrating data from two primary sources (**Methods** and **Supplementary Table 1**): (1) publicly available human gut metagenomic datasets comprising 9,320 samples from 39 previous studies and (2) 3,584 newly sequenced fecal samples from healthy Han Chinese. All these samples are from 22 countries, including 5,631 samples from 29 representative provinces and regions in China (**Figure 1a** and **Supplementary Table 1**). We used our in-house metagenomic assembly workflow (**Methods**) to generate 210,269 MAGs for downstream analyses. Additionally, we collected 85,801 microbial genomes (**Supplementary Table 2**) from established human gut genome catalogs [11, 13–15, 19, 21] (**Methods**).

**Fig. 1.**
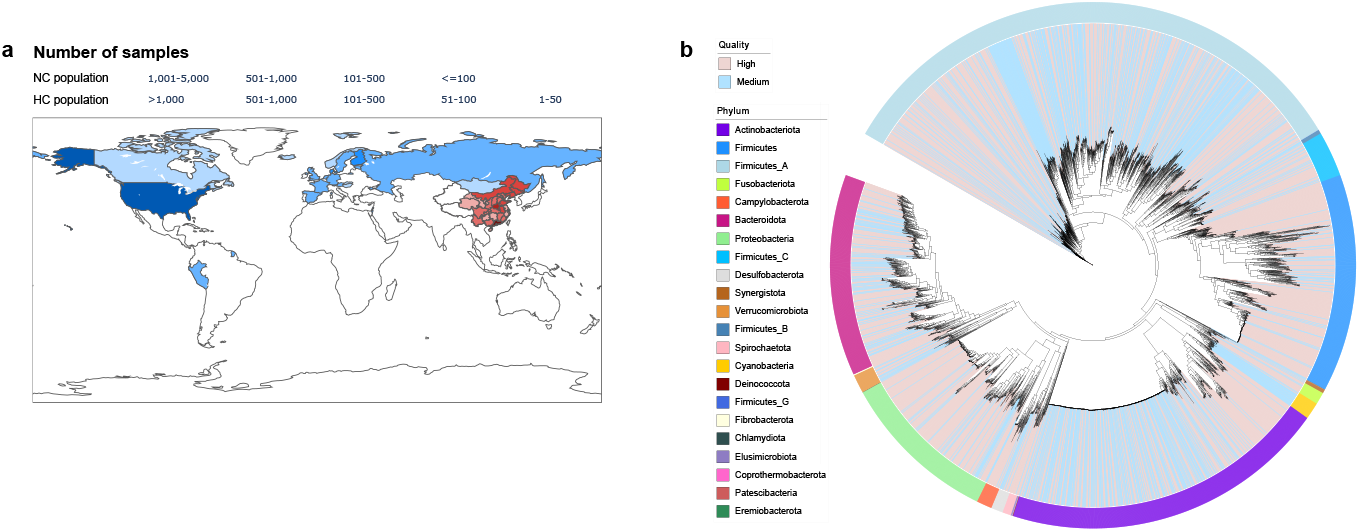
A high-quality human gut microbial reference genome catalog. **a** Geographic distribution of metagenomic samples collected from 22 countries and 29 provinces or regions across mainland China. **b** Maximum-likelihood phylogenetic tree comprising 5,740 representative genomes from bacteria. Clades are colored based on genome quality (high-quality genomes (completeness ≥ 90% and contamination ≤ 5%) in pink, medium-quality genomes (completeness ≥ 75%, contamination ≤ 10%, and QS ≥ 50) in blue), and the outside circle shows the phylum annotation.

After genome clustering and dereplication (**Methods**), we generated 5,785 species-level genome clusters including 271,480 non-redundant genomes (**Supplementary Table 3**). We selected a representative genome (RG) for each cluster (**Methods**), revealing a total of 5,740 (99.22%) RGs from bacteria (**Figure 1b**) and 45 (0.78%) RGs from archaea distributed across 26 distinct phyla (**Supplementary Table 4**). Detailed descriptive statistics of HGMRGC can be found in the **Supplementary Notes** and **Supplementary Figure 1**.

Compared to the UHGG, HGMRGC included 66,542 more non-redundant genomes and 1,141 more species-level genome clusters. The RGs in HGMRGC also exhibited higher completeness, lower contamination, larger N50 and longer genome length (**Supplementary Figure 2**). Moreover, HGMRGC demonstrated a 19.87% improvement in phylogenetic diversity and a higher mapping rate in external metagenomic datasets (**Supplementary Table S5**) than UHGG. Detailed comparative analysis of HGMRGC and UHGG can be found in the **Supplementary Notes** and **Supplementary Figure 2**. Our reference genome catalog provided a high-quality and diverse human gut microbial genome collection to support microbial SNPs detection and downstream analysis.

### Geography as the primary driver of gut microbial variation

We constructed the discovery and replication cohorts to further explore the difference in gut microbiota between the HC and NC populations (**Methods**). The discovery cohort comprised 2,762 HC and 1,680 NC individuals sourced from previous studies (**Supplementary Table 6**). The replication cohorts included 2,449 HC individuals sequenced by this study and 601 NC individuals from TwinsUK [22] (**Supplementary Table 6**). Our analysis (**Methods**) revealed geography as the most influential factor driving microbial variation in both abundance and SNP-based tests, surpassing age, sex and BMI (**Figure 2a-b**). Geography explained the largest proportion of microbial variation, accounting for 3.43% (3.89%) of abundance-based variation and 2.55% (1.94%) of SNP-based variation in the discovery (replication) cohorts, respectively (PERMANOVA test, *p* < 0.0001; **Figure 2a-2b**). Age was the second most influential factor, significantly contributing to microbial variation among individuals (PERMANOVA test, *p* < 0.0001; **Figure 2a-2b**).

**Fig. 2.**
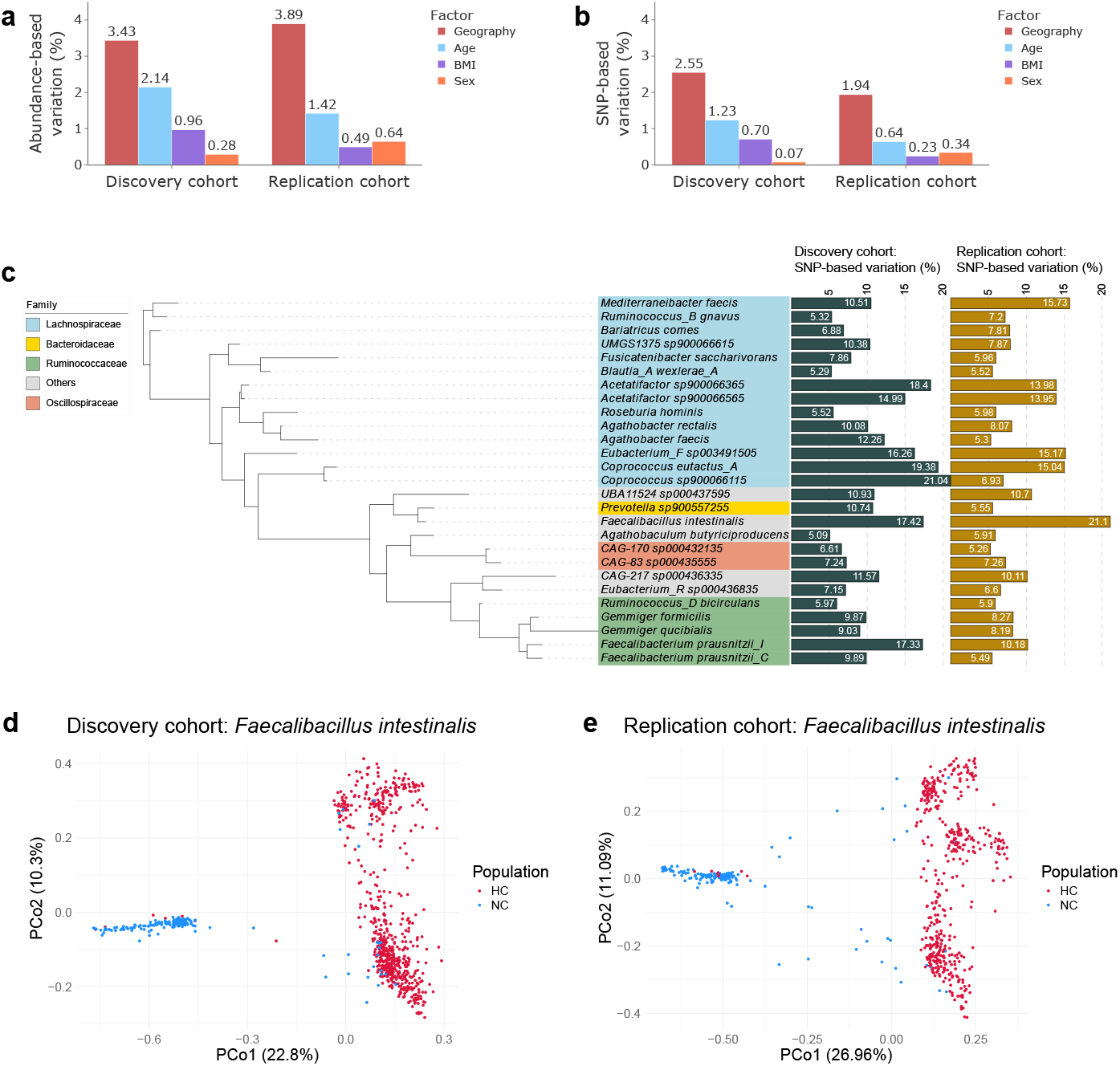
The contributions of geography, age, sex, BMI to microbial variation among individuals. **a-b** The variation explained by geography, age, sex and BMI based on microbial abundance (**a**) and SNP (**b**) in the discovery and replication cohorts using PERMANOVA test (*p* < 0.0001). **c** Maximum-likelihood phylogenetic tree for 27 genomes that geography can explain more variation (≥ 5%) than the other factors (PERMANOVA test, *p* < 0.0001). The background colors of tree node labels represent their annotated families. **d-e** Principal coordinates analysis of the differences in *Faecalibacillus intestinalis* between HC and NC populations based on SNP-based microbial profiles in the discovery (**d**) and replication (**e**) cohorts. The percentage of variation explained by the first and second principal coordinates (PCo1 and PCo2) was indicated on the axes.

We also examined the SNP-based test on microbial variation for each RG (**Methods**). In both the discovery and replication cohorts, we identified 27 RGs (**Supplementary Table 7**) for which the geography factor accounted for at least 5% of SNP-based inter-individual variation and surpassed the proportions of variation explained by age, sex and BMI (**Figure 2c**). We found more than half (14 of 27) of these RGs belonged to the Lachnospiraceae family. The metabolic phenotype analysis (**Methods**) revealed two categories of phenotypes, amino acid and nucleotide biosynthesis, as mostly conserved across the Lachnospiraceae family (**Supplementary Figure 3** and **Supplementary Table 8**). In contrast, carbohydrate utilization and vitamin synthesis exhibited the greatest variability among different RGs of this family (**Supplementary Figure 3** and **Supplementary Table 8**). The fermentation-related phenotypes, such as acetate, formate and butyrate production, also played critical functional roles in RGs of the Lachnospiraceae family (**Supplementary Figure 3** and **Supplementary Table 8**). We observed *Faecalibacillus intestinalis*, a bacterium known to be influenced by dietary interventions in healthy human faces [23], exhibited the strongest geographical influence, with geography explaining 17.42% and 21.10% of its SNP-based inter-individual variation in the discovery and replication cohorts, respectively (**Figure 2c**). Furthermore, the principal coordinates analysis (PCoA) (**Methods**) revealed a clear geographical pattern in the SNP-based variation of *Faecalibacillus intestinalis* between the HC and NC populations (**Figure 2d-2e**). The first principal coordinate (PCo1) accounted for over 20% (**Figure 2d-2e**) of the total SNP-based variation, highlighting its dominant role in differentiating the two populations. Strikingly, 85.96% (81.13%) of NC samples exhibited PCo1 values below −0.4, whereas 94.13% (98.43%) of HC samples displayed PCo1 values above 0 in the discovery (replication) cohort. To investigate the functional implications of this population-level divergence, we examined the SNP-based alpha diversity of genes in *Faecalibacillus intestinalis* between NC and HC individuals (**Methods**). Thirty-four genes (**Supplementary Table 9**) showed significantly different SNP-based alpha diversity in both cohorts (two-tailed Mann–Whitney U test, *p* < 0.05), including six linked to carbohydrate and lipid metabolism KEGG modules. For example, the gene *fabD*, encoding malonyl-CoA-acyl carrier protein transacylase, was directly involved in fatty acid biosynthesis (M00082).

### Beta diversity and selection pressure of gut microbiota between populations

To assess microbial community divergence, we calculated abundance-based and SNP-based pairwise beta diversity using Bray-Curtis dissimilarity and allele-sharing scores (**Methods**). Our analyses demonstrated significantly higher beta diversity between populations compared to within populations in both the discovery and replication cohorts (two tailed Mann–Whitney U test, *p* < 0.0001; **Figure 3a-3b**). We further applied the fixation index (*F*_*ST*_) to assess selection pressures on individual RGs between populations (**Methods**). The *F*_*ST*_ values of different RGs ranged from 0.69 to 0.99, highlighting substantial variability in selection pressures among different microbial species (**Supplementary Table 10** and **Figure 3c**). Seventeen of the top 20 RGs (85%) exhibiting the highest *F*_*ST*_ values in the discovery cohort were validated in the independent replication cohort. These included six genomes from the Lachnospiraceae family, four from the Acutalibacteraceae family, three from the Bacteroidaceae family, and four from other families (**Figure 3c**). To investigate functional selection signals, we calculated *F*_*ST*_ for protein-coding genes in RGs (**Methods**). We observed that 19 of the top 20 genes ranked by *F*_*ST*_ in the discovery cohort also exhibited robust selection signals in the replication cohort (**Supplementary Table 11**). Among these 19 genes, we identified two genes related to carbohydrate metabolism: *bglB* (encoding phospho-beta-glucosidase B) from *Bacteroides uniformis*, and *xynB* (encoding xylanase) from *Phocaeicola dorei*. These enzymes play key roles in digesting dietary fibers and plant-derived compounds, facilitating the production of short-chain fatty acids that are beneficial to gut health. Additionally, we observed the gene *rarA* with high *F*_*ST*_ values in *Bacteroides uniformis*, which is associated with antibiotic resistance and bacterial survival under antibiotic pressure [24].

**Fig. 3.**
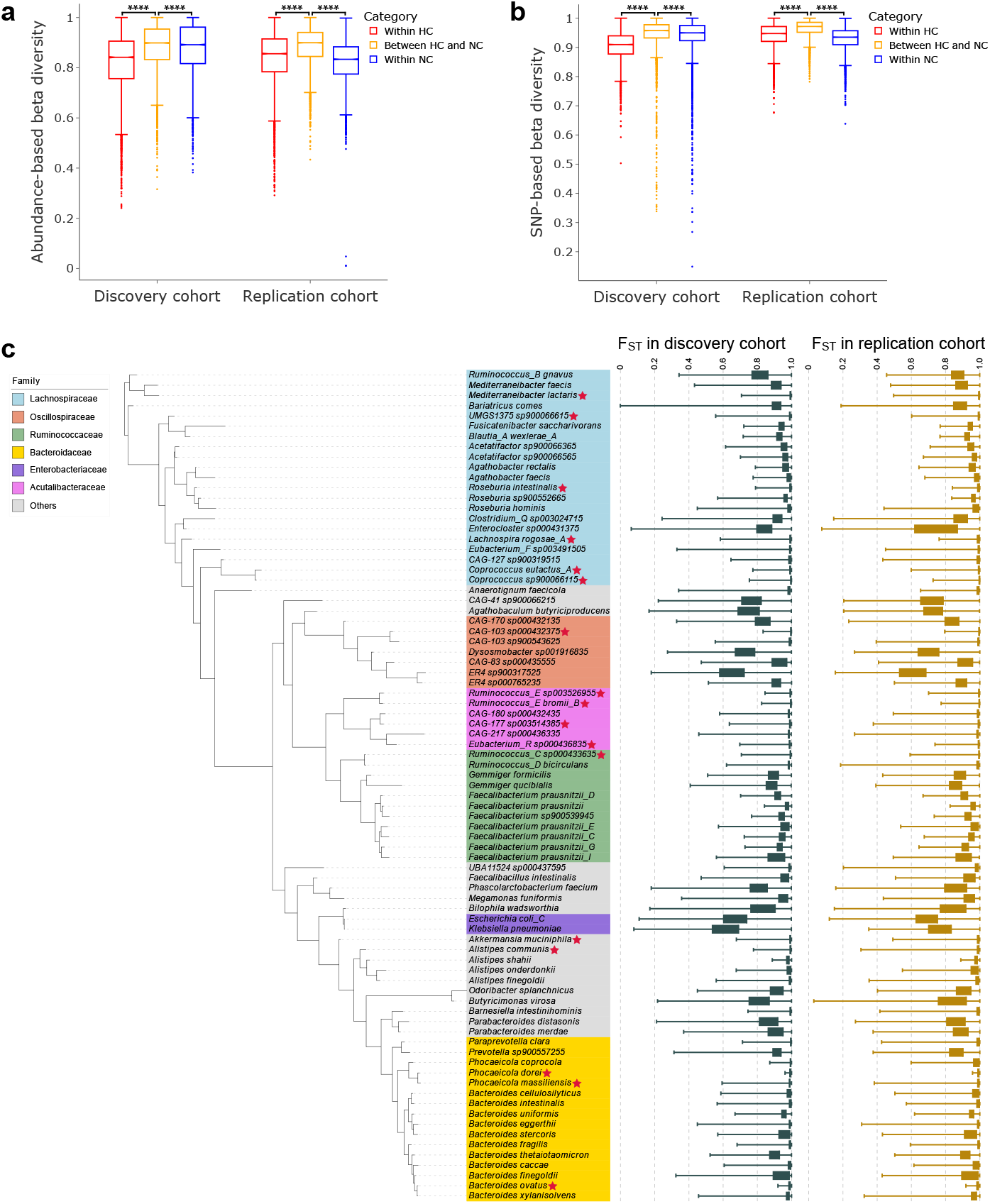
Beta diversity and selection pressure between HC and NC populations. **a-b** The abundance-based beta diversity (**a**) and SNP-based beta diversity (**b**) within HC population (red), within NC population (blue) and between HC and NC populations (yellow). The values of abundance- and SNP-based beta diversity were derived from 10,000 sample pairs that were randomly selected from each of the three groups. Two-tailed Mann–Whitney U tests assessed the differences between groups, **** *p<* 0.0001. **(c)** Maximum-likelihood phylogenetic tree comprising the 81 eligible representative genomes for the analysis of selection pressure between populations. The two box plots show the *F*_*ST*_ distributions of genomes in the discovery and replication cohorts. The red stars highlight the 17 genomes that ranked among the top 20 genomes with the highest average *F*_*ST*_ values in the discovery cohort and were also replicated in the replication cohort. The background colors of tree node labels represent their annotated families.

### Population-specific novel species highlight microbial differences in carbohydrate utilization

We identified 625 novel genome clusters (**Supplementary Table 12**) in HGMRGC, all of which were derived exclusively from metagenome assemblies using sequencing data collected or generated by this study and were not reported in the previous research. These novel genome clusters demonstrated unique population-specific characteristics, including 356 (56.95%) and 256 (40.96%) clusters where all involved genomes were only from the HC and NC populations, respectively. Interestingly, we observed that 99 and 139 population-specific novel RGs (PSNRGs) (**Supplementary Table 12**) from the HC and NC populations were allocated in two separate branches of *Collinsella* genus in the phylogenetic tree (**Figure 4a**), which suggested that there might be some functional differences of these RGs between the two populations.

**Fig. 4.**
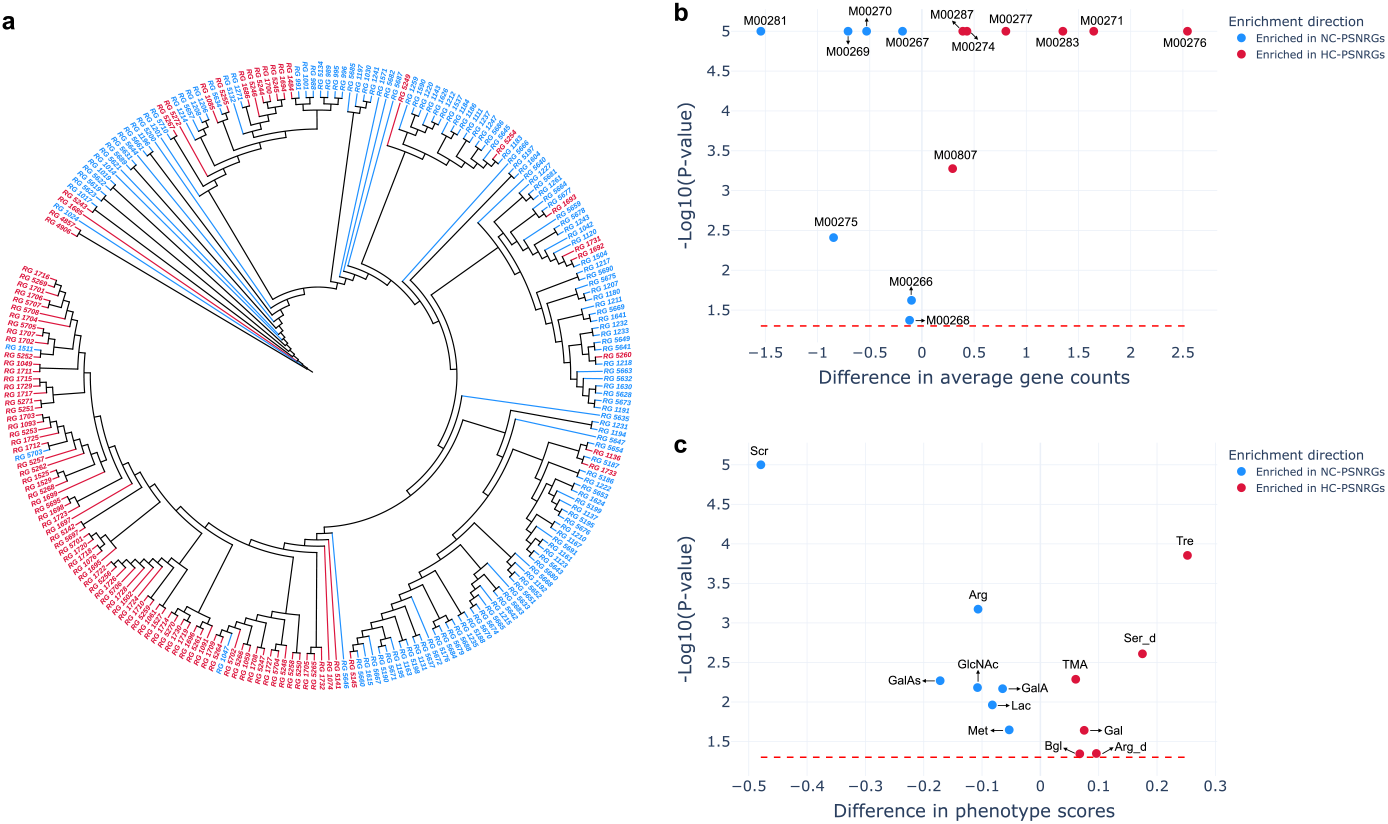
Population-specific novel representative genomes from the *Collinsella* genus. **a** Maximum-likelihood phylogenetic tree comprising 238 novel representative genomes from the *Collinsella* genus. HC-specific representative genomes (99) are in red and NC-specific representative genomes (139) are in blue. **b** Scatter plot shows the enrichment analysis of KEGG modules between HC-PSNRGs and NC-PSNRGs. X-axis indicates the difference in the average number of genes involved in the KEGG module between HC-PSNRGs and NC-PSNRGs. Y-axis indicates the log10 transformation of the *p* values generated from one-tailed permutation tests (permutation times = 100,000). A dashed red line marks the *p* value threshold of 0.05. Each point on the plot represents a KEGG module, colored by enrichment direction: red for the KEGG module enriched in HC-PSNRGs and blue for the KEGG module enriched in NC-PSNRGs. Text near each point indicates the KEGG module ID, “M00271” represents the beta-glucoside PTS module, “M00281” represents the lactose PTS module, detailed description of other modules can be found in the **Supplementary Table 13. c** Scatter plot shows the enrichment analysis of metabolic phenotypes between HC-PSNRGs and NC-PSNRGs. X-axis indicates the difference in phenotype scores between HC-PSNRGs and NC-PSNRGs. Y-axis indicates the log10 transformation of the *p* values generated from one-tailed permutation tests (permutation times = 100,000). A dashed red line marks the *p* value threshold of 0.05. Each point on the plot represents a metabolic phenotype, colored by enrichment direction: red for the phenotype enriched in HC-PSNRGs and blue for the phenotype enriched in NC-PSNRGs. Text near each point indicates the phenotype, “Lac” represents lactose utilization, “Bgl” represents “beta-glucoside utilization”. Detailed description of other phenotypes can be found in the **Supplementary Table 13**.

We annotated protein-coding genes for the 238 PSNRGs from the *Collinsella* genus and compared the average number of genes across KEGG pathways and modules between PSNRGs in the HC (HCPSNRGs) and NC (NC-PSNRGs) populations (**Methods**). Significant differences were observed in the phosphotransferase system (PTS) (map02060) (one-tailed permutation test, *p* < 0.0001), a critical carbohydrate transport system essential for sugar utilization and energy production [25]. Specifically, we observed different distributions of genes in PTS-related KEGG modules between the HC-PSNRGs and NC-PSNRGs (**Supplementary Table 13**). For example, HC-PSNRGs have significantly more genes involved in the beta-glucoside PTS module (M00271) (mean: 4.49 vs. 2.85, one-tailed permutation test, *p* < 0.0001; **Figure 4b**), whereas the number of genes involved in the lactose PTS module (M00281) was significantly higher in the NC-PSNRGs (mean: 1.41 vs.2.96, one-tailed permutation test, *p* < 0.0001; **Figure 4b**). Metabolic phenotypes of *Collinsella* PSNRGs further highlighted these functional disparities (**Supplementary Table 13**). HC-PSNRGs demonstrated substantially enrichment for beta-glucoside utilization (phenotype score: 0.96 vs. 0.89 in NC-PSNRGs; one-tailed permutation test, *p* < 0.0001; **Figure 4c**), whereas NC-PSNRGs were strongly associated with lactose utilization (phenotype score: 0.97 vs. 0.89 in HC-PSNRGs; one-tailed permutation test, *p* < 0.0001; **Figure 4c**). These results align genomic signatures with metabolic capabilities, highlighting niche-specific adaptations in carbohydrate metabolism.

### Differences in microbial prevalence between populations

We identified 126 population-specific prevalent RGs (PSPRGs) (**Supplementary Table 14**) that had significant prevalence differences between the two populations in both the discovery and replication cohorts (**Methods**). There were 62 and 64 PSPRGs in the HC (HC-PSPRGs) and NC (NC-PSPRGs) populations, respectively (**Figure 5a**). The PSPRGs were distributed across 39 different families, with over half (58%) of them belonging to five dominant families: Lachnospiraceae (27%), Oscillospiraceae (11%), Ruminococcaceae (8%), Bacteroidaceae (7%) and Enterobacteriaceae (5%) (**Figure 5b**). Notably, the Bacteroidaceae and Enterobacteriaceae families showed distinct population-specific characteristics: almost all PSPRGs within these two families were HC-PSPRGs (Bacteroidaceae: 8 of 9, Enterobacteriaceae: 6 of 6). In contrast, NC-PSPRGs were enriched in the Ruminococcaceae family (7 of 10) (**Figure 5b**).

**Fig. 5.**
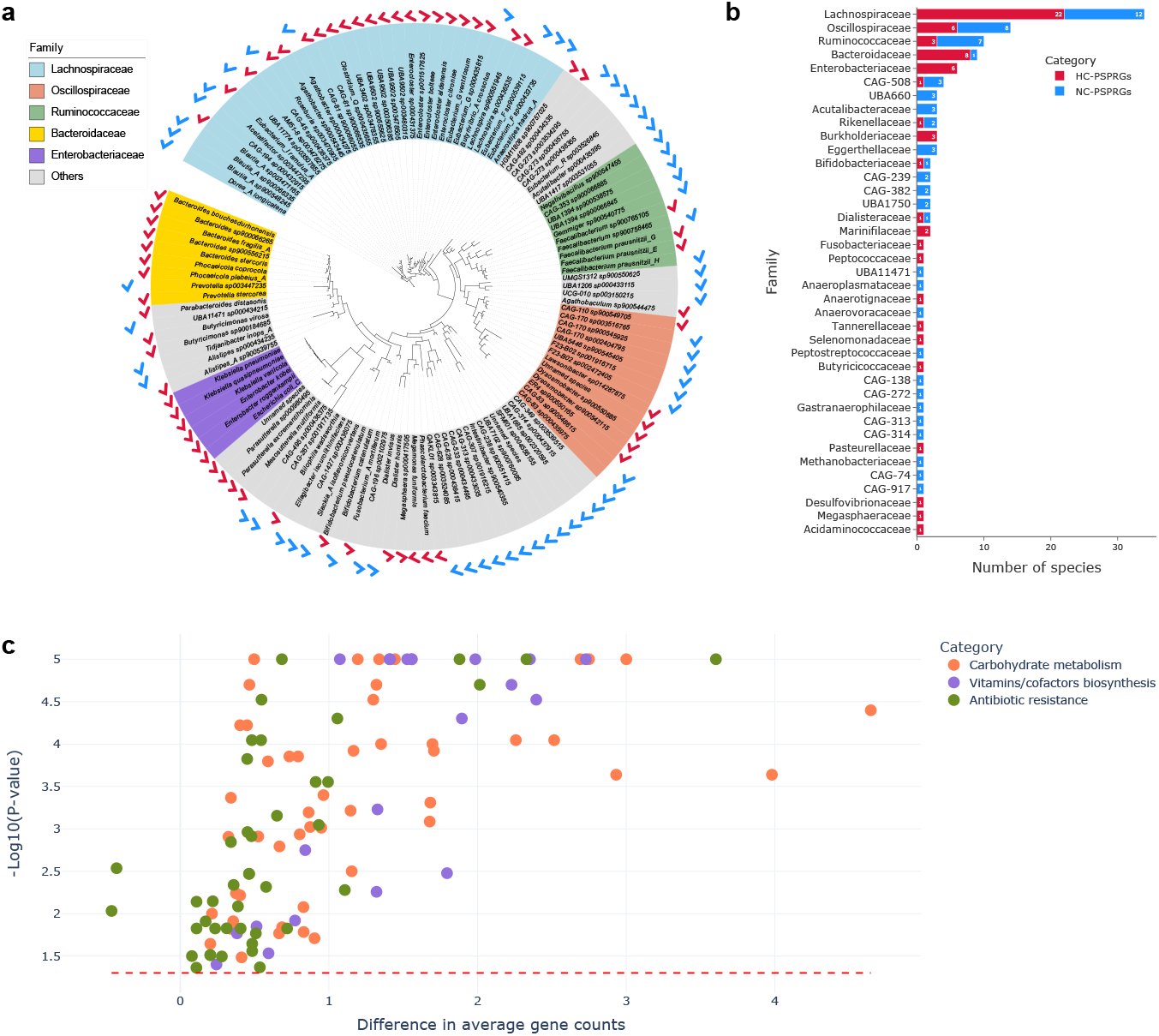
Microbial prevalence differences between HC and NC populations. **a** Maximum-likelihood phylogenetic tree comprising 126 population-specific prevalent genomes. Red and blue checkmarks annotate the 62 HC-PSPRGs and 64 NC-PSPRGs, respectively. The background colors of tree node labels represent their annotated families. **b** The distribution of annotated families for the 126 population-specific prevalent genomes. **c** Scatter plot shows the enrichment analysis of KEGG modules between HC-PSPRGs and NC-PSPRGs. X-axis indicates the difference in the average number of genes involved in the KEGG module between HC-PSPRGs and NC-PSPRGs. Y-axis indicates the log10 transformation of the *p* values generated from one-tailed permutation tests (permutation times = 100,000). A dashed red line marks the *p* value threshold of 0.05. Each point on the plot represents a KEGG module, colored by KEGG module category: carbohydrate metabolism in orange, vitamins/cofactors biosynthesis in purple, antibiotic resistance in green. Detailed description of KEGG modules can be found in the **Supplementary Table 15**.

Our functional enrichment analysis (**Methods**) revealed that the HC-PSPRGs were enriched in the functions associated with carbohydrate metabolism, antibiotic resistance and vitamins biosynthesis (**Figure 5c** and **Supplementary Table 15**). HC-PSPRGs demonstrated significantly higher enrichment for the glucose utilization metabolic phenotype compared to NC-PSPRGs (phenotype score: 0.56 vs. 0.31, one-tailed permutation test, *p* < 0.01) and exhibited a higher gene count associated with the glycolysis KEGG module (M00001: Embden-Meyerhof pathway), a core glucose-to-pyruvate conversion pathway (mean: 0.11 vs. 0.03, one-tailed permutation test, *p* = 0.02). HC-PSPRGs carried over twice as many genes associated with the cationic antimicrobial peptide (CAMP) resistance pathway (map01503) compared to NC-PSPRGs (mean: 12.42 vs. 5.35, one-tailed permutation test, *p* < 0.0001). Similarly, genes linked to beta-lactam resistance (map01501) showed a nearly 1.5-fold increase in HC-PSPRGs relative to NC-PSPRGs (mean: 15.98 vs. 10.85, one-tailed permutation test, *p* = 0.002). HC-PSPRGs contained significantly more genes involved in the biosynthesis of B vitamins (including B1, B2, B5, B6, B7, B9, and B12) and vitamin K2 compared to NC-PSPRGs (**Supplementary Table 15**). This trend was further validated by the metabolic phenotype analysis, which revealed significant enrichment of the phenotypes linked to *de novo* synthesis of these vitamins in HC-PSPRGs (**Supplementary Table 15**), highlighting their strong metabolic capacity for vitamin production. Additionally, HC-PSPRGs also demonstrated significant enrichment for the trimethylamine (TMA) production phenotype (phenotype score: 0.16 vs. 0, one-tailed permutation test, *p* < 0.001), which is strongly associated with cardiovascular diseases [26].

### Alpha diversity of gut microbiota within populations

We observed significant differences in microbial alpha diversity within the two populations (**Figure 6a-6b**). Compared to the HC population, the NC population exhibited significantly higher abundance-based alpha diversity (**Methods**) in both the discovery (median: 4.50 for HC, 5.13 for NC, two-tailed Mann–Whitney U test, *p* < 0.0001; **Figure 6a**) and replication cohorts (median: 4.67 for HC, 5.40 for NC, two-tailed Mann–Whitney U test, *p* < 0.0001; **Figure 6a**). These findings align with previous studies indicating that the HC population typically has lower gut microbial diversity compared to other ethnic groups with respect to microbial abundance [27–29]. In contrast, the SNP-based alpha diversity (**Methods**) demonstrated an opposing trend, with the HC population showing significantly higher diversity than the NC population (**Figure 6b**). This pattern was consistent across the discovery (median: 0.0578 for HC, 0.0424 for NC, two-tailed Mann–Whitney U test, *p* < 0.0001) and replication cohorts (median: 0.0585 for HC, 0.0558 for NC; two-tailed Mann–Whitney U test, *p* < 0.0001), even the NC population in the discovery cohort comprising individuals from multiple countries (**Supplementary Table 6** and **Methods**).

**Fig. 6.**
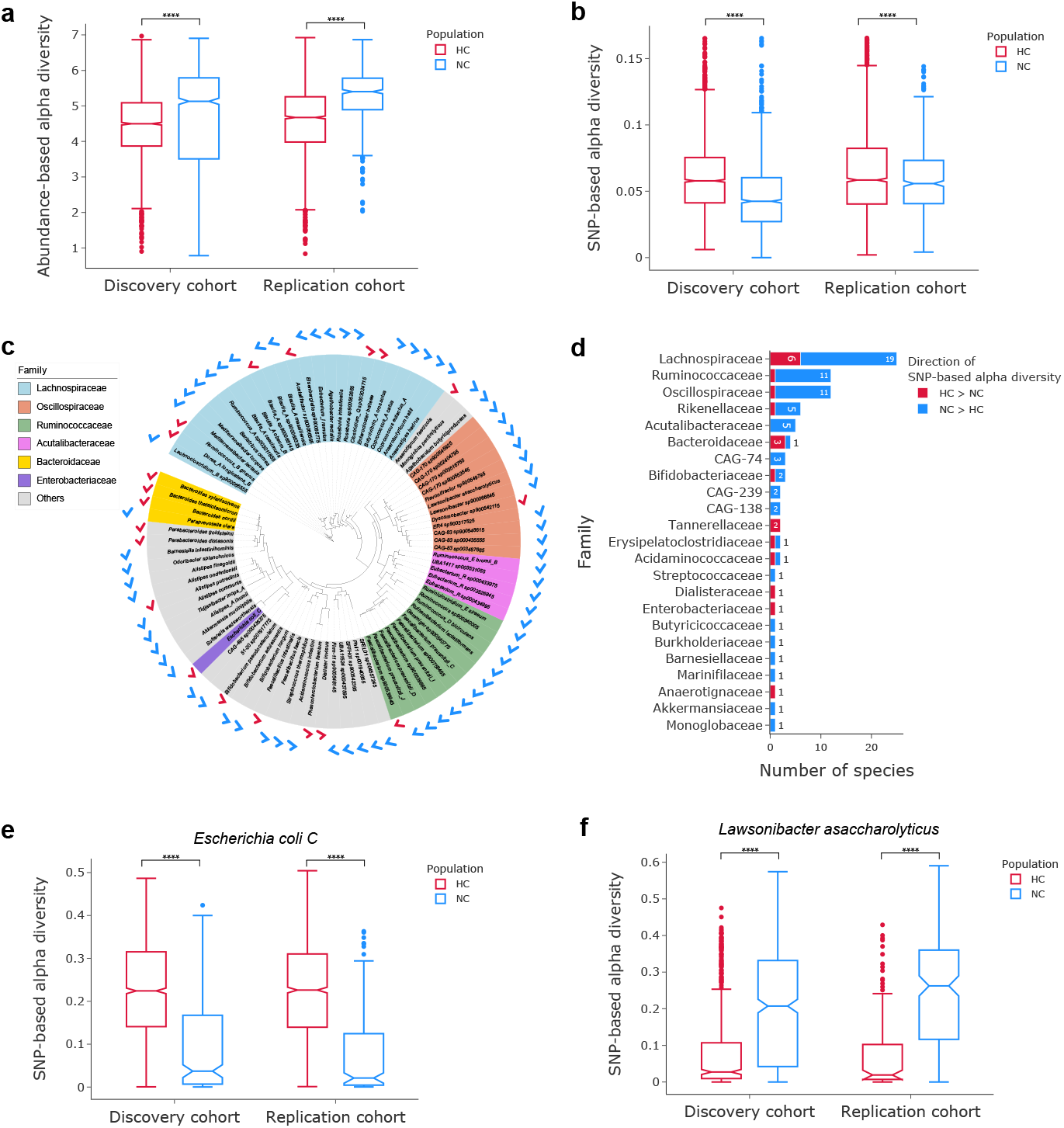
Alpha diversity differences between HC and NC populations. **a-b** Box plots of the abundance-based alpha diversity (**a**) and the SNP-based alpha diversity (**b**) in the discovery and replication cohorts. Two tailed Mann–Whitney U tests assessed the differences between populations, **** *p* < 0.0001. **c** Maximum-likelihood phylogenetic tree comprising the 90 representative genomes with significant differences in SNP-based alpha diversity in both two cohorts. The red (blue) checkmarks annotate 20 (90) representative genomes with higher SNP-based alpha diversity in HC (NC) population. The background colors of tree node labels represent their annotated families. **d** The distribution of annotated families for the 90 representative genomes with significant differences in SNP-based alpha diversity. **e** SNP-based alpha diversity of *Escherichia coli C* in the discovery (HC: n=1,788; NC: n=284) and replication (HC: n=1,653; NC: n=243) cohorts. Two-tailed Mann–Whitney U tests assessed the differences between the two populations, **** *p* < 0.0001. **f** SNP-based alpha diversity of *Lawsonibacter asaccharolyticus* in the discovery (HC: n=491; NC (n=603) and replication (HC: n=129; NC: n=193) cohorts. Two-tailed Mann–Whitney U tests assessed the differences between the two populations, **** *p<* 0.0001.

We further examined the SNP-based alpha diversity for individual RGs to identify single bacteria with significant differences between populations (**Methods**). Notably, our analysis revealed that 20 genomes demonstrated significantly higher alpha diversity in the HC population compared to 70 genomes in the NC population in both the discovery and replication cohorts (**Figure 6c** and **Supplementary Table 16**). These genomes spanned 23 families, with three dominant families-Lachnospiraceae (28%), Oscillospiraceae (13%), and Ruminococcaceae (13%)-accounting for over half (54%) of the significant genomes (**Figure 6d**). For Oscillospiraceae and Ruminococcaceae, almost all significant genomes in these two families displayed higher SNP-based alpha diversity in the NC population (Oscillospiraceae: 11/12, Ruminococcaceae: 11/12) (**Figure 6d**).

We observed significant differences in the SNP-based alpha diversity of *Escherichia coli C* between the two populations in both the discovery (median: 0.2249 for HC, 0.0923 for NC, two-tailed Mann– Whitney U test, *p* < 0.0001; **Figure 6e**) and replication cohorts (median: 0.2204 for HC, 0.0768 for NC; two-tailed Mann–Whitney U test, *p* < 0.0001; **Figure 6e**). *Escherichia coli C* facilitates lactose metabolism in the human gut by producing beta-galactosidase, which hydrolyzes lactose into glucose and galactose [30]. Three lactose metabolism-related genes in the *Escherichia coli C* genome were identified: *lacZ* (encoding beta-galactosidase), *lacY* (encoding lactose permease for lactose transport), and *lacA* (encoding galactoside O-acetyltransferase). All three genes exhibited significantly lower SNP-based alpha diversity in the NC population. Additionally, *Lawsonibacter asaccharolyticus*, an intestinal bacteria associated with coffee consumption [31], was identified with substantially higher SNP-based alpha diversity in the NC population, with mean diversity more than twice that of the HC population in both the discovery (median: 0.0869 for HC, 0.2073 for NC, two-tailed Mann–Whitney U test, *p* < 0.0001; **Figure 6f**) and replication (median: 0.0755 for HC, 0.2484 for NC, two-tailed Mann–Whitney U test, *p* < 0.0001; **Figure 6f**) cohorts.

## Discussion

The human gut microbiota, often referred to as the “second genome” [1], shaped by a complex interplay of host genetics, diet, and environmental factors and demonstrates substantial differences between populations [32]. In this study, we expanded the scope of analysis to examine the effects of microbial SNPs by utilizing our high-quality and diverse human gut microbial genome collection HGMRGC. Our findings revealed that geography is the most influential factor shaping both composition and SNP in the human gut microbiota, surpassing other factors such as age, sex and BMI. We also systematically compared the gut microbiota between the HC and NC populations, with respect to population-specific species derived from metagenome assemblies, microbial prevalence and diversity based on both abundance and SNP data. The observed divergence in gut microbiome between the two populations likely reflects the significant dietary distinctions, vitamin synthesis and antibiotic resistance.

The traditional dietary patterns of Han Chinese populations, historically characterized by minimal lactose consumption due to limited dairy intake, is in contrast to Western dietary practices that emphasize frequent milk and dairy product consumption [33]. This divergence in nutritional exposure may drive microbial evolutionary adaptations, as evidenced by the substantial enrichment of lactose utilization phenotypes within NC-PSNRGs from the *Collinsella* genus. Such genomic signatures likely reflect selection pressures imposed by dairy-rich diets, highlighting the role of cultural dietary habits in shaping functional gut microbiome profiles. Besides the *Collinsella* genus, we found that *Escherichia coli C* in NC populations showed significantly reduced SNP-based alpha diversity in lactose metabolism-associated genes (*lacZ, lacY*, and *lacA*). This low genetic variability suggests that these genes may be under stronger evolutionary selection or functional conservation in the NC groups compared to HC populations. We found *Faecalibacillus intestinalis* demonstrated a substantial geographical pattern between the two populations, with significant differences in SNP-based alpha diversity of genes related to carbohydrate and lipid metabolism. The bacterium *Faecalibacillus intestinalis* has been reported to be highly responsive to dietary interventions, with evidence suggesting that low-fat, low-sugar, and high-fiber diets could significantly influence its abundance in the human gut microbiota [34]. We also observed the species from Lachnospiraceae family exhibited significant differences between HC and NC populations, which have been recognized to play a critical role in digesting complex carbohydrates and producing short-chain fatty acids [35–37]. The *bglB* gene in *Bacteroides uniformis*, responsible for encoding phospho-beta-glucosidase B, is under significant selective pressure and plays a pivotal role in the metabolism of dietary complex carbohydrates and plant fibers. The enrichment of beta-glucoside utilization and related PTS module in HC-PSNRGs further indicates a greater capacity for HC individuals to metabolize complex carbohydrates and plant fibers. In our study, we further identified a significant difference in the SNP-based alpha diversity of *Lawsonibacter asaccharolyticus* between the HC and NC populations, which has been identified to be associated with coffee consumption [31]. This difference may be partially attributable to differences in coffee consumption patterns, as such dietary practice is more prevalent in NC populations.

Differences in gut microbiota between the HC and NC populations were also evident in vitamin synthesis and TMA production. HC-PSPRGs showed significant enrichment of the *de novo* synthesis for vitamins B and K2. Vitamin K2 participates in cardiovascular protection by activating coagulation factors and maintaining vascular elasticity [38], while vitamins B6, B9, and B12 can regulate homocysteine metabolism and reduce the level of blood homocysteine [39]. An elevated level of homocysteine is recognized as a risk factor for cardiovascular diseases [40]. Previous research reported that gut bacteria-driven production of TMA is strongly associated with cardiovascular disease, potentially exacerbating its progression [26]. However, the TMA-producing metabolic phenotype was enriched in HC-PSPRGs, but absent in NC-PSPRGs. These findings suggest that the gut microbiota in HC populations may influence cardiovascular health through a “double-edged sword” metabolic regulation: on one hand, enhanced vitamins B and K2 synthesis capabilities may confer protective effects through homocysteine clearance and coagulation homeostasis; on the other hand, the active TMA production pathway could counteract these protective effects. This dual influence may partially explain the higher risk of cardiovascular diseases in the Chinese population compared to those in the USA and European countries [41].

The prevalence analysis revealed significant enrichment of antibiotic resistance in HC populations, such as CAMP and beta-lactam resistance. This finding aligned with previous studies indicating that the Chinese gut microbiome harbors a higher number of antibiotic resistance genes compared to other populations [42, 43]. Antibiotic usage was another important driver of population-specific microbial characteristics [44]. In China, widespread and often excessive antibiotic use has contributed to elevated levels of antibiotic resistance genes in gut microbiota [45], whereas Western countries’ stricter regulations have mitigated such effects [46].

This study highlights the importance of considering both abundance-based and SNP-based diversity to comprehensively investigate changes in microbial communities. In our study, the beta diversity analysis highlighted significant inter-population differences in both microbial abundance and SNPs, while alpha diversity analysis showed NC populations exhibited higher diversity in microbial abundance, and HC populations showed greater diversity in microbial SNPs. These findings suggest that relying solely on abundance-based diversity metric may not fully capture the complexity of microbial community structures. The difference in abundance-based alpha diversity could be linked to dietary habits, environmental factors, and lifestyle [27–29], whereas the SNP-based alpha diversity may reflect evolutionary selection in single bacterial species. Future research should explore the ecological and evolutionary implications of these diversity differences and assess their potential impact on host health. Besides exploring the differences between HC and NC populations, we also identified significant substructures of the human gut microbiota within populations. The PCoA analysis on *Faecalibacillus intestinalis* revealed two subclusters within HC populations along the second principal coordinate (PCo2) : one comprising HC individuals with PCo2 values below 0 (HC-subcluster 1) and the other with PCo2 values above 0.2 (HC-subcluster 2), observed in both the discovery and replication cohorts (**Figure 2d-2e**). We found genes such as *metE* and *metH* from *Faecalibacillus intestinalis*, involved in methionine biosynthesis (M00017), with significant higher SNP-based alpha diversity in HC-subcluster 2 (**Supplementary Table 17**). These findings indicate the necessity of considering fine-scale within population structure in the future.

## Methods

### Data Collection and metagenomic sequencing

We collected short-read metagenomic sequencing data from 9,320 human fecal samples available in the European Nucleotide Archive (ENA) [47], sourced from 39 distinct studies (**Supplementary Table 1**). Geographical metadata for each sample was retrieved from the ENA or supplementary materials of the respective publications. Additionally, we generated metagenomic sequencing data from 3,584 samples collected from the Han Chinese population. These samples were obtained from healthy volunteers across 26 provinces and regions in mainland China, all of whom provided written informed consent. The study received approval from the Ethics Committee of Guangdong Provincial Hospital of Chinese Medicine (Ethics Batch Number: B2017-199-01, Date of Approval: 2017-12-15). Volunteers who had taken antibiotics, gastrointestinal motility drugs, or gut microbiota-affecting supplements within the previous three months were excluded. The fecal samples of these volunteers were collected using stool collection kits (MCK-01 KMHD, Shenzhen, China) and stored at −80°C. Genomic DNA (minimum 3 *µ*g) was extracted using the QIAsymphony PowerFecal Pro DNA kit (QIAGEN, Germany). Shotgun sequencing libraries were prepared with the TruSeq DNA HT Sample Prep Kit (Illumina, USA) and sequenced on the HiSeq X Ten and NovaSeq platforms to produce 150bp paired-end reads. The adapter contamination and low-quality reads were filtered out using Kneaddata v.0.12.1 (https://github.com/biobakery/kneaddata), and any reads were removed if they could be aligned to the human reference genome (GRCh38) using Bowtie2 v.2.5.4 [48].

### Discovery and Replication cohorts

The discovery cohort consisted of 2,762 HC and 1,680 NC individuals sourced from 11 previous studies (**Supplementary Table 5**). This cohort included 51% female and 49% male, with an average age of 40.24 years (s.d. = 16.14) and an average BMI of 24.27 (s.d. = 6.03). The replication cohort consisted of 2,449 HC individuals sequenced by this study and 601 NC individuals from the TwinsUK study [22]. This cohort included 52% female and 48% male, with an average age of 51.11 years (s.d. = 13.55) and an average BMI of 24.43 (s.d. = 3.79). The NC individuals in the discovery cohort represented 11 different countries, while those in the replication cohort were all from the UK.

### Metagenome assembly and Contig binning

Raw reads from metagenomic sequencing were assembled into contigs using metaSPAdes v.3.15.3 [49]. We performed contig binning using MetaBAT 2 v.2.12.1 [50] with a minimum contig length threshold of 1,500 bp.

### Extract microbial genomes from RefSeq and available reference genome catalogs

Unbinned contigs from our assemblies were mapped to the complete bacterial genomes in the human gut from the RefSeq collection [21] using minimap2 v2.26-r1175 [51]. Genomes were included in our catalog if unbinned contigs covered at least 60% of their sequences. Additionally, 6,284 human gut isolate genomes [10–12, 21] and 79,457 genomes [11, 13–15, 19] derived from previous studies were included into our catalog.

### MAG genome quality evaluation

We used CheckM v1.1.2 [52] to evaluate the completeness and contamination of MAGs. The quality score (QS) for each genome was calculated as completeness − 5×contamination. HGMRGC only kept high-quality MAGs (completeness ≥ 90% and contamination ≤ 5%) and medium-quality MAGs (completeness ≥ 75%, contamination ≤ 10%, and QS ≥ 50) for subsequent analyses.

### Microbial genome clustering and dereplication

We grouped all the collected microbial genomes into three categories: high-quality genomes, medium-quality genomes, and complete genomes (assembled from bacterial isolation or the qualified complete genomes from the RefSeq). For each category, we applied hierarchical clustering at a Mash distance (Mash v2.0 [53]) threshold of 0.2 to generate preliminary genome clusters. Each genome cluster was further dereplicated using dRep v2.5.4 [54]. In each cluster, we selected the genome with the highest genome score, calculated by dRep v2.5.4 [54], as a RG. RGs from medium-quality genome clusters were aligned with those from high-quality genome clusters using the mashdiff.sh [13]. It calculated the average nucleotide identity (ANI) and aligned query (AQ) values for each pair of RGs using Mash v2.0 [53] and dnadiff v1.3 from MUMmer v3.23 [55]. Medium-quality and high-quality genome clusters were merged if their RGs satisfied an AQ *>* 60% and an ANI *>* 95%. The RG from the original high-quality genome cluster was selected as the RG of the merged cluster. This procedure was repeatedly applied by aligning RGs from high-quality, medium-quality, and previously merged genome clusters with those from complete genome clusters. To further alleviate redundancy in each cluster, we dereplicated all conspecific genomes at a 99.9% ANI threshold using dRep v2.5.4 [54].

### Comparison of HGMRGC and UHGG

We used one-tailed Mann–Whitney U test to compare genome completeness, contamination, N50 and genome length between HGMRGC and UHGG. We aligned additional metagenomic sequencing data from 200 human fecal samples (**Supplementary Table 14**) to RGs in the HGMRGC and UHGG by BWA-MEM 0.7.17-r1188 [56] to compare their read mapping rates. These samples included 100 HC individuals from the CNGB Sequence Archive (accession number CNP0000334), and 100 NC individuals from the TwinsUK cohort [22]. The number of mapped and unmapped reads was extracted from the statistics file generated by the “SAMtools stats” [57]. The mapping rate was calculated as the ratio of mapped reads to the total reads. The significance of read mapping rate differences was evaluated by one-sided Wilcoxon signed-rank test.

### Taxonomic annotation and phylogenetic analysis

Taxonomic annotation of 5,785 RGs in HGMRGC was conducted using GTDB-Tk v1.5.0 (database r202) [58] with the “classify_wf” function. GTDB-Tk [58] also identified universal marker genes for each RG. We concatenated the marker genes for the 5,740 bacterial RGs, and a bacterial phylogenetic tree was constructed using IQ-TREE v2.1.4beta [59]. Tree visualization and annotation were performed using iTOL [60]. Phylogenetic diversity was quantified as the sum of branch lengths using the phytools package [61].

### Gene prediction and functional annotation

The protein-coding sequences (CDSs) for each RG were predicted using Prokka v1.14.6 [62]. The functions of all CDSs, including their involved KEGG modules and KEGG pathways, were annotated using eggNOG-mapper v2.1.11 [63] with the database v5.0.2 [64].

### Metabolic phenotype annotation

To annotate the metabolic phenotypes of 5,785 RGs in HGMRGC, we performed our in-house phenotype prediction workflow based on the mcSEED database [65]. This database includes a genomics-based collection of 158 reconstructed metabolic pathways for key nutrients and intermediary metabolites, derived from 3,441 reference bacterial genomes that represent the human gut microbiota [65]. The presence or absence of each of the 158 functional metabolic pathways (binary phenotypes) in RGs were identified based on the mcSEED database, thereby generating a Binary Phenotype Matrix (BPM) as previously described [66]. The BPM reflects the inferred presence or absence of 158 metabolic phenotypes in each analyzed RG.

### Calculate microbial abundances and prevalence

We initially aligned metagenomic sequencing reads to RGs using BWA-MEM 0.7.17-r1188 [56] and removed poorly aligned reads by SAMtools v.1.6 (SAMtools view -q 1 -rf 2) [57]. We calculated Reads Per Kilobase of transcript per Million mapped reads (RPKM) to quantify the relative abundance for each RG. A genome was considered present in the population only if it met two criteria: (1) genome coverage exceeding 40%, and (2) a RPKM value greater than 0.0001.

### Detection of SNPs for metagenomic sequencing data

We selected 922 species-level genome clusters from HGMRGC, each containing at least 10 high-quality genomes (completeness ≥ 90% and contamination ≤ 5%). To ensure reliable and efficient SNP calling, 64 species-level genome clusters with more than 1,000 conspecific genomes were downsampled to 1,000 conspecific genomes based on their genome quality scores. We performed whole-genome alignment between all pairs of conspecific genomes in the same cluster using MUMmer v4.0.0 [67]. Based on pairwise genome similarity, we identified the template genome as the one with the highest average similarity to all others. A total of 59.3 million common, core-genome bi-allelic SNPs (minor allele frequency ≥ 1%, site prevalence ≥ 90%) were identified by aligning each conspecific genome to the template genome using Masst [68]. Leveraging this extensive SNP catalog of the human gut microbiome, we created a SNP-covering kmer (sck-mers, k=31) database containing 5.9 billion unique species-specific sck-mers using GT-Pro [69]. The SNPs for each sample were identified by matching k-mers of short-reads from the sample with the sck-mer database using GT-Pro [69]. To filter out false positive SNPs from GT-Pro outputs, we excluded SNPs from the species that were not present in the sample. Additionally, we further removed SNPs with read depth less than 4 and alternative allele frequency below 0.01.

### Alpha diversity calculation

We evaluated the abundance-based alpha diversity using the Shannon index calculated by QIIME 2 [70]. A single-site nucleotide diversity (*π*) was assessed using the formula [69]:

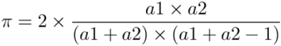

where *ai* (*i* = 1 or 2) represents the coverage of the *i*th genotype of a bi-allelic SNP site. The SNP-based alpha diversity for a metagenomic sequencing sample was calculated as the average *π* value across all SNP sites within the sample. Similarly, the SNP-based alpha diversity for RGs and genes was determined by averaging *π* values across all SNP sites within the respective regions.

### Beta diversity calculation

We applied the Bray-Curtis dissimilarity as the abundance-based beta diversity to quantify the species compositional distance between two samples. The Bray-Curtis dissimilarity between samples was calculated using QIIME 2 [70]. To calculate the SNP-based beta diversity, we first determined the allele-sharing score [69] between each pair of samples based on their SNP genotype results. The SNP-based beta diversity between two samples was then calculated as 1 - allele-sharing score for those samples.

### Interpretation of microbial variation

We conducted a permutational multivariate analysis of variance (PERMANOVA) to assess the contributions of different factors to microbial abundance-based and SNP-based variation. PERMANOVA is a non-parametric method that evaluates the significance of differences between groups based on a distance matrix. To construct the abundance-based and SNP-based distance matrices, we calculated the respective beta diversity among individuals. The PERMANOVA incorporated four explanatory variables, including geography, age, sex, and BMI. Specifically, the geographical variable indicated whether an individual originated from mainland China. PERMANOVA test was performed using the “permanova” function from the scikit-bio library (http://scikit-bio.org) with 100,000 times permutation.

### Principal coordinates analysis

We performed PCoA based on the SNP-based beta diversity matrix of *Faecalibacillus intestinalis*. The classical multidimensional scaling was performed using the “cmscale” function from the vegan [71] to calculate different principal coordinates. The results of the PCoA were visualized using scatter plots generated with the ggplot2 [72].

### *F*_*ST*_ **calculation**

To calculate *F*_*ST*_ for RGs, we randomly selected 6,000 sample pairs, each consisting of one sample from the HC population and one from the NC population. This analysis included 81 RGs meeting the following criteria: (1) a prevalence exceeding 10% in both populations, and (2) the presence of over 1,000 shared SNP sites across the sample pairs. The *F*_*ST*_ was calculated as [69]:

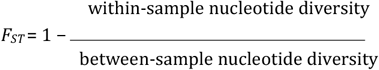

between-sample nucleotide diversity. To estimate within-sample nucleotide diversity, we first calculated single-site nucleotide diversity (*π*) and then averaged these values across all SNP sites [69]. For between-sample nucleotide diversity, we computed single-site nucleotide diversity (*π*_*b*_) for shared SNP sites, and subsequently averaged *π*_*b*_ across all shared SNP sites [69]. For each sample pair, *F*_*ST*_ was calculated, and the average *F*_*ST*_ value across all pairs was used to represent the genome-level *F*_*ST*_ between populations. Similar method was used to calculate the gene-level *F*_*ST*_ between populations.

### Functional analysis for HC-PSNRGs and NC-PSNRGs

For each KEGG module, we calculated the average gene count per genome involved in this module between the 99 HC-PSNRGs and 139 NC-PSNRGs. A one-tailed permutation test was conducted to assess the significance of the difference in average gene counts between HC-PSNRGs and NC-PSNRGs (*p* < 0.05). For each permutation, 99 and 139 genomes were randomly sampled from the total set of 238 PSNRGs to compute the difference between average gene counts. This procedure was repeated 100,000 times to create an empirical distribution of the test statistic. Similar permutation tests were performed to compare the difference in KEGG pathways between HC-PSNRGs and NC-PSNRGs. For the metabolic phenotype analysis, we calculated the phenotype score, defined as the frequency of the metabolic phenotype in HC-PSNRGs or NC-PSNRGs. Similar permutation tests were performed to assess the significance of the difference in the phenotype score between the HC-PSNRGs and NC-PSNRGs.

### Identify HC-PSPRGs and NC-PSPRGs

We employed two-tailed Fisher’s exact test to identify species with significantly different prevalence between the HC and NC populations (*p* < 0.05). The statistical analysis was performed using the “fisher_exac” function in the SciPy package [73]. For the significant species, we applied two criteria to define HC-PSPRGs (NC-PSPRGs): (1) the prevalence in the NC (HC) population must be below 5%, and (2) the prevalence in the HC (NC) population must be at least three times higher than in the NC (HC) population.

### Functional analysis for HC-PSPRGs and NC-PSPRGs

For each KEGG module, we calculated the average gene count per genome involved in this module between the 62 HC-PSPRGs and 64 NC-PSPRGs. A one-tailed permutation test was conducted to assess the significance of the difference in average gene counts between HC-PSPRGs and NC-PSPRGs (*p* < 0.05). For each permutation, 62 and 64 genomes were randomly sampled from the total set of 126 PSPRGs to compute the difference between average gene counts. This procedure was repeated 100,000 times to create an empirical distribution of the test statistic. Similar permutation tests were performed to compare the difference in KEGG pathways between HC-PSPRGs and NC-PSPRGs. For the metabolic phenotype analysis, we calculated the phenotype score, defined as the frequency of the metabolic phenotype in HC-PSPRGs or NC-PSPRGs. Similar permutation tests were performed to assess the significance of the difference in the phenotype score between the HC-PSPRGs and NC-PSPRGs.

### Statistical analysis for alpha and beta diversity

The comparison of alpha diversity between the HC and NC populations was conducted using two complementary statistical analyses in both the discovery and replication cohorts. First, a two-sided Mann–Whitney U test was conducted to assess whether the distributions of alpha diversity differed significantly between the two populations. Second, a one-tailed permutation test was conducted to determine the direction of the observed differences by comparing the differences in average alpha diversity between the two populations. Statistical analyses were performed using the “mannwhitneyu” and “permutation test” functions from the SciPy package [73]. Each permutation was repeated 100,000 times, and the result with a significance level of *p* < 0.05 in both tests were selected for further analysis. A similar statistical approach was employed to compare the beta diversity between the two populations. Beta diversity values were calculated from 10,000 randomly selected sample pairs within each of the three groups: between HC and NC populations, within the HC population, and within the NC population.

## Acknowledgements

We thank all the participants for their significant contribution to this study. The design of the study and the collection, analysis and interpretation of the data were supported by the Young Collaborative Research grant (No. C2004-23Y), HMRF grant (No. 11221026), the State Key Laboratory of Dampness Syndrome of Chinese Medicine (No. SZ2024KF22), National Natural Science Foundation of China (No. 82004234), Science and Technology Planning Project of Guangdong Province, China (No. 2020B1111100005).

## Data availability

The 9,320 public-available metagenomic sequencing data can be accessed in the European Nucleotide Archive under 39 different accession numbers (**Supplementary Table 1**). The 3,584 metagenomic sequencing data generated in this study are available in the CNGB Sequence Archive under the accession number CNP0006798. The metagenomic sequencing data from 4,442 individuals in the discovery cohort were obtained from 11 projects (**Supplementary Table 9**). The metagenomic sequencing data from 2,449 HC individuals can be accessed in the CNGB Sequence Archive under the accession number CNP0006798. The metagenomic sequencing data from 601 NC individuals in the replication cohort were sourced from the TwinsUK study [22] and can be accessed by submitting a request through the TwinsUK website (http://twinsuk.ac.uk/resources-for-researchers/access-our-data/). By accessing the HGMRGC web website (http://www.hgmrgc.com:4380/downloads), users can download the 5,785 representative genomes, their annotations, taxonomy and genome statistics.

## Code availability

Commands and parameters used to create our reference genome catalog are available in the **Supplementary Notes**. The source code for prediction of metabolic phenotype presence or absence in RGs is available at https://github.com/rodionovdima/PhenotypePredictor-2 and has been accessioned at Zenodo (DOI: 10.5281/zenodo.13250651).

## Author contributions

L.Z and X.D.F conceived the study. Y.C, Q.W.Q, Y.S.D, Y.M.L, C.R.W, X.X.S, L.H, C.S, J.W.G, Z.M.Y, L.J.H, and L.X.Z contributed to the acquisition of the metagenomic sequencing data. Z.M.Z performed the genome assembly, clustering and dereplication. J.J.W annotated genomes, profiled metagenomic sequencing reads, conducted the analyses, interpreted the results, and drafted the manuscript. D.A.R and A.L.O performed the metabolic phenotype annotation. J.J.W and X.Z, and D.A.R pre-preprocessed and analyzed the metagenome profiling data. J.J.W, Y.C, and C.Y analyzed and interpreted the results. J.X.X developed the website. L.Z, X.D.F, W.J, Z.X.B, J.J.W, Z.M.Z, and Y.C revised the manuscript. All authors had access to the study data, reviewed and approved the final manuscript.

