## Supplementary Notes for "Exploring Population Differences in the Human Gut Microbiome: from Microbial Abundance to Single Nucleotide Polymorphisms"

### Descriptive statistics of HGMRGC

HGMRGC was achieved by using metagenomic sequencing data from 9,320 publicly available samples across 22 countries and 3,584 newly sequenced samples from the Han Chinese individuals across 29 provinces and regions **(Supplementary Table 1)**. The average sequencing amount for each sample was 6.60 Gb for Han Chinese (HC) and 4.05 Gb for non-Chinese (NC) populations (**Supplementary Table 1**). There were 5,785 prokaryotic species-level genome clusters including 271,480 non-redundant genomes (**Supplementary Table 3**) in HGMRGC. We selected a representative genome (RG) for each cluster (**Methods**), resulting in a total of 4,757 (82.2%) high-quality (completeness ≥ 90%, contamination ≤ 5%) and 1,028 (17.8%) medium-quality (completeness ≥ 75%, contamination ≤ 10%, quality score ≥ 50) RGs (**Supplementary Figure 1a** and **Supplementary Table 4**).

We utilized the Genome Taxonomy Database Toolkit (GTDB-Tk V1.5.0, database R202) to assign taxonomic lineages for all 5,785 RGs (**Methods**), which revealed 5,740 (99.22%) RGs from bacteria and 45 (0.78%) RGs from archaea, respectively (**Supplementary Table 4**). A total of 4,505 (77.9%) RGs could be annotated at the species level, while 1,189 (20.6%) and 91 (1.5%) RGs were assigned to the genus and above the genus levels, respectively (**Supplementary Figure 2a**). All of these RGs were distributed across 23 bacterial and 3 archaeal phyla (**Supplementary Table 4**). The five predominant phyla were Firmicutes A, Firmicutes, Bacteroidetes, Actinobacteria, and Proteobacteria, which collectively comprised 90.39% of the total RGs (**Supplementary Figure 1b**). Notably, over 90% of the RGs from Firmicutes and Proteobacteria phyla were high-quality (**Supplementary Figure 1c**). The phylum Firmicutes A showed the greatest phylogenetic diversity (271.75), which was more than two times of the second largest phylum Firmicutes (103.89) (**Supplementary Figure 1d**). Through metabolic phenotype annotation (**Methods**), we observed that the RGs within the five predominant phyla were primarily linked to metabolic phenotypes involving carbohydrate utilization and amino acid synthesis (**Supplementary Figure 1e**).

### Comparative analysis of HGMRGC and UHGG

Compared to the UHGG, HGMRGC included 66,542 more non-redundant genomes (HGMRGC: 271,480, UHGG: 204,938) and 1,141 more representative genomes (HGMRGC: 5,785, UHGG: 4,644). In the UHGG, there were 88.44% (4,107 of 4,644) (**Supplementary Table 2**) representative genomes that satisfied our stringent MAG quality criteria of completeness (**Methods**). Comparing representative genomes between the two catalogs, our representative genomes exhibited higher completeness (median completeness: 97.66%[HGMRGC], 95.69%[UHGG], one-tailed Mann–Whitney U test, $p < 0.0001$; **Supplementary Figure 2b**), lower contamination (median contamination: 0.60%[HGMRGC], 0.67%[UHGG], one-tailed Mann–Whitney U test, $p = 0.002$; **Supplementary Figure 2c**), higher N50 (median N50: 75.95 Kbp[HGMRGC], 48.50 Kbp[UHGG], one-tailed Mann–Whitney U test, $p < 0.0001$; **Supplementary Figure 2d**), larger genome length (median genome length: 2.25 Mbp[HGMRGC], 2.14 Mbp[UHGG], one-tailed Mann–Whitney U test, $p < 0.0001$; **Supplementary Figure 2e**), and more average non-redundant genomes per clusters (46.93[HGMRGC], 44.13[UHGG]). We also found that HGMRGC showed a 19.87% improvement in phylogenetic diversity (**Methods**) across all phyla compared to the UHGG (HGMRGC: 625.58, UHGG: 522.29). HGMRGC has 974 more genomes being annotated at the species level than that of the UHGG (HGMRGC: 4,505, UHGG: 3,531; **Supplementary Figure 2a**). We randomly selected 200 external metagenomic samples (100 HC samples and 100 NC samples, **Supplementary Table 5**) and aligned short-reads to the reference genomes from the HGMRGC and UHGG (**Methods**). We observed the mapping rate was significantly improved using the HGMRGC (median mapping rate: 95.56%[HGMRGC], 95.08%[UHGG], one-tailed Wilcoxon signed-rank test, $p < 0.0001$; **Supplementary Figure 2f**).

### Legends for Supplementary Tables

**Supplementary Table 1: Overview of the human gut metagenomic sequencing data collection.** This table summarizes the collection of the human gut metagenomic sequencing data analyzed in this study. The dataset includes publicly available metagenomic sequencing data from 9,320 human fecal samples, sourced from 39 independent studies and retrieved from the European Nucleotide Archive. Geographic metadata (continent, country, and province) for each sample were obtained from the ENA records or supplementary materials of the corresponding publications. Additionally, this study contributed newly sequenced metagenomic data from 3,584 Han Chinese fecal samples, collected from healthy volunteers across 26 provinces and regions in mainland China.

**Supplementary Table 2: Summary of gut genomes utilized in this study.** This table provides an overview of the genomes utilized in this study, including genomes collected from existing human gut genome catalogs and metagenome-assembled genomes (MAGs) generated in this work. We grouped all the collected microbial genomes into three categories: high-quality genomes (completeness ≥ 90%, contamination ≤ 10%), medium-quality (completeness ≥ 75%, contamination ≤ 10%, quality score ≥ 50) genomes, and complete genomes (assembled from bacterial isolation or the qualified complete genomes from RefSeq).

**Supplementary Table 3: List of the 271,480 non-redundant genomes grouped into 5,785 species-level genome clusters.** This table presents the clustering of 271,480 non-redundant genomes into 5,785 species-level genome clusters. Genomes sharing the same 'Genome Cluster ID' belong to the same species-level cluster. The column 'Representative or Not' indicates whether the genome serves as a representative genome within its respective cluster.

**Supplementary Table 4: List of the 5,785 representative genomes in HGMRGC.** This table provides the list of representative genomes included in the Human Gut Microbial Reference Genome Collection (HGMRGC). The taxonomic classification for each representative genome was assigned using GTDB-Tk v1.5.0 (database r202).

**Supplementary Table 5: Mapping rates of the metagenomic sequencing data from 200 external individuals.** This table presents the mapping rates for metagenomic sequencing data from 200 external individuals, including 100 Han Chinese (HC) samples from the CNGB Nucleotide Sequence Archive (accession number CNP0000334) and 100 non-Chinese (NC) samples from the TwinsUK cohort. The mapping rate was calculated as the proportion of reads mapped to the reference catalog (HGMRGC or UHGG) relative to the total number of reads.

**Supplementary Table 6: Individual metadata of the discovery and replication cohorts.** This table shows the individual metadata of the discovery and replication cohorts. The discovery cohort includes 2,762 Han Chinese (HC) and 1,680 non-Chinese (NC) individuals from 11 previous studies. The replication cohort comprises 2,449 HC individuals sequenced in this study and 601 NC individuals from the TwinsUK study.

**Supplementary Table 7: List of the 27 representative genomes with significant SNP-based variation attributed to geography.** This table lists the 27 representative genomes for which geographical factors account for at least 5% of the SNP-based inter-individual variation, surpassing the variation contributions from age, sex, and BMI in both discovery and replication cohorts. The estimated variation (%) was assessed using the PERMANOVA method ($p$< 0.0001).

**Supplementary Table 8: Metabolic phenotypes of the 14 representative genomes in the Lachnospiraceae family.** This table lists the metabolic phenotypes of the 14 Lachnospiraceae representative genomes with significant SNP-based variation attributed to geography.

**Supplementary Table 9: List of genes with significant difference in SNP-based alpha diversity for *Faecalibacillus intestinalis* between HC and NC populations.** This table lists the 34 genes in *Faecalibacillus intestinalis* that exhibit significantly different SNP-based alpha diversity between NC and HC individuals.

**Supplementary Table 10: $F_{ST}$ values of representative genomes between populations.** This table shows $F_{ST}$ values of representative genomes between the HC and NC populations. For each eligible reference genomes, we randomly selected 6,000 sample pairs for $F_{ST}$ calculation, each consisting of one sample from the HC population and one from the NC population. The average $F_{ST}$ value across all pairs was used to represent the genome-level $F_{ST}$ between populations.

**Supplementary Table 11: $F_{ST}$ values of genes from representative genomes between populations.** This table shows $F_{ST}$ values of genes from representative genomes between the HC and NC populations. For each gene, we randomly selected 6,000 sample pairs for $F_{ST}$ calculation, each consisting of one sample from the HC population and one from the NC population. The average $F_{ST}$ value across all pairs was used to represent the gene-level $F_{ST}$ between populations.

**Supplementary Table 12: List of the 625 novel genome clusters in HGMRGC.** This table provides the list of 625 novel genome clusters identified in HGMRGC. All genomes within the novel clusters were derived exclusively from MAGs using sequencing data collected or generated by this study and were not reported in previous research. The novel genome clusters are categorized based on their population origins: HC population-specific clusters include genomes exclusively from the HC population, NC population-specific clusters include genomes exclusively from the NC population, and mixed population-specific clusters include genomes from both the HC and NC populations.

**Supplementary Table 13: List of the KEGG modules, pathways and metabolic phenotypes with significantly different differences between HCPSNRGs and NC-PSNRGs from the *Collinsella* genus.** This table displays the KEGG modules, pathways and metabolic phenotypes with significantly different differences between HC-PSNRGs and NC-PSNRGs from the *Collinsella* genus, along with detailed descriptions of related KEGG modules, pathways and metabolic phenotypes.

**Supplementary Table 14: List of the 126 PSPRGs.** This table lists the 126 PSPRGs that had significant prevalence differences between the two populations in both discovery and replication cohorts.

**Supplementary Table 15: List of the KEGG modules, pathways and metabolic phenotypes with significantly different differences between HCPSPRGs and NC-PSPRGs.** This table displays the KEGG modules, pathways and metabolic phenotypes with significantly different differences between HC-PSPRGs and NC-PSPRGs, along with detailed descriptions of related KEGG modules, pathways and metabolic phenotypes.

**Supplementary Table 16: List of the 90 representative genomes with significant differences in SNP-based alpha diversity between populations.** This table lists the 90 representative genomes with significant differences in SNP-based alpha diversity between populations.

**Supplementary Table 17: List of genes with significant difference in SNP-based alpha diversity for *Faecalibacillus intestinalis* between HC-subcluster 1 and HC-subcluster 2.** This table lists the genes in *Faecalibacillus intestinalis* that exhibit significantly different SNP-based alpha diversity between HC-subcluster 1 and HC-subcluster 2.

### Commands and parameters used to construct and annotate the HGMRGC

- **Contig assembly**

  Metaspades.py -o metaspades_out -1 R1.fastq -2 R2.fastq

- **Contig binning**

  metabat2 -m 1500 -assembly.fasta -o outdir

- **Genome dereplication**
  - For medium-quality genomes

    dRep dereplicate output_directory -g genomes.fasta -pa 0.9 -sa 0.95 -cm larger -nc 0.30
  - For high-quality genomes

    dRep dereplicate output_directory -g genomes.fasta -pa 0.9 -sa 0.95 -cm larger -nc 0.60
  - For complete genomes

    dRep dereplicate output_directory -g genomes.fasta -pa 0.9 -sa 0.95 -cm larger -nc 0.60
  - Remove redundant genomes

    dRep dereplicate output_directory -g genomes.fasta -pa 0.999 --SkipSecondary

- **Functional annotation**
  - Prokka for bacterial genomes

    prokka --outdir mydir --prefix mygenome contigs.fa --kingdom Bacteria
  - Prokka for archaeal genomes

    prokka --outdir mydir --prefix mygenome contigs.fa --kingdom Archaea
  - EggNOG-mapper

    emapper.py --override -i prokka_output.faa –output out_prefix –output_dir out_dir --excel

- **Taxonomic annotation**

  gtdbtk classify_wf --batchfile batch.txt -out_dir our_dir

- **Phylogenetic tree construction**

  iqtree -s dup_msa_bac.fasta
