## Supplementary Figures for "Exploring Population Differences in the Human Gut Microbiome: from Microbial Abundance to Single Nucleotide Polymorphisms"

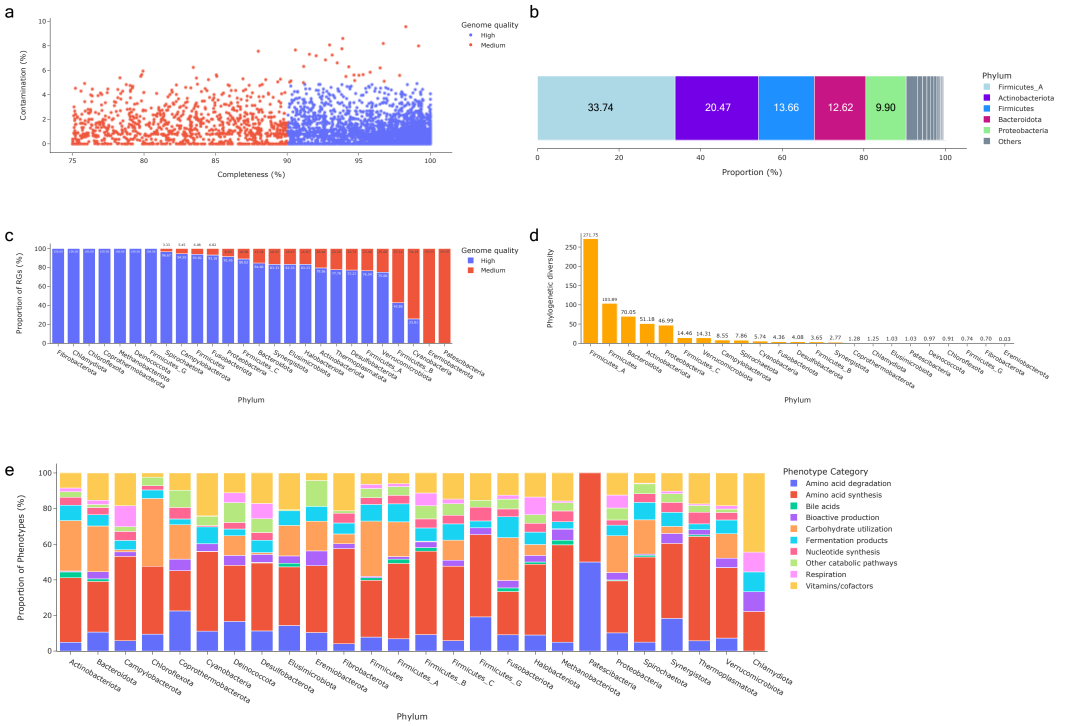


**Fig. S1 Descriptive statistics of HGMRGC. a** Completeness and contamination scores for each of the 5,785 representative genomes, colored by their quality classification category. High-quality: completeness ≥ 90%, contamination ≤ 10%; medium-quality: completeness ≥ 75%, contamination ≤ 10%, quality score ≥ 50. Quality score = completeness − 5 × contamination. **b** The proportion of representative genomes in each phylum. **c** The proportion of high-quality and medium-quality representative genomes in each phylum. **d** The phylogenetic diversity of each phylum. **e** The proportion of metabolic phenotypes for representative genomes in each phylum.

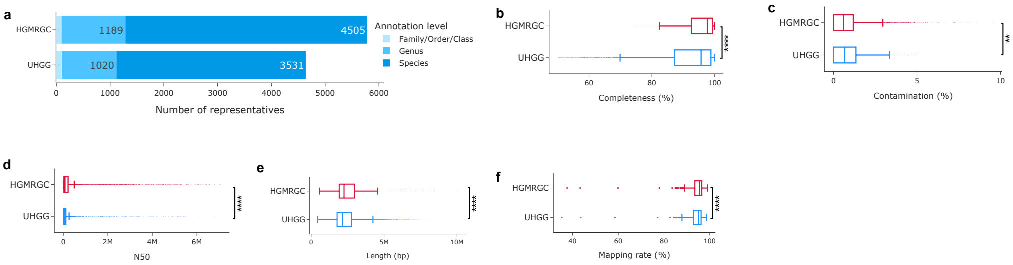


**Fig. S2 Comparison of representative genomes in HGMRGC and UHGG. a** The number of representative genomes annotated at the lowest taxonomic levels in HGMRGC and UHGG. **b-e** Box plots of the completeness (**b**), contamination (**c**), N50 (**d**), length of the representative genomes (**e**). HGMRGC (n=5,785) in red and UHGG (n=4,644) in blue. One-tailed Mann–Whitney U test assessed the differences between HGMRGC and UHGG, **** $p < 0.0001$, ** $p < 0.01$. **f** Box plot shows the read mapping rates for 200 external metagenomic sequencing samples. One-tailed Wilcoxon signed-rank test assessed the differences between HGMRGC and UHGG, **** $p < 0.0001$.

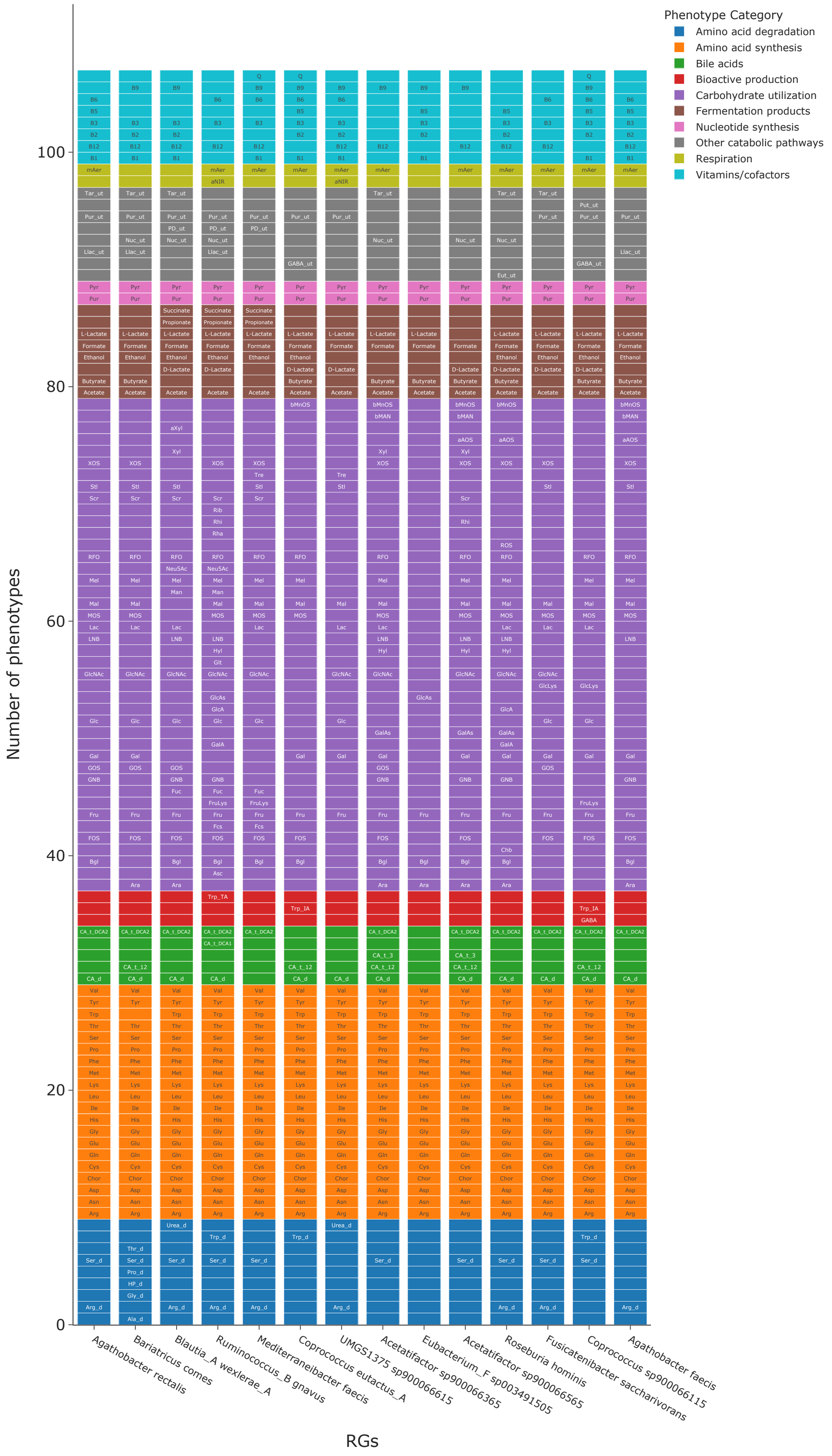


**Fig. S3 Metabolic phenotypes of the 14 representative genomes from the Lachnospiraceae family.** Text within the cell indicates phenotype abbreviation, which is present only if the corresponding phenotype is present in the RG. Detailed description of each metabolic phenotype can be found in the **Supplementary Table 8**.
